# Single cell based high-throughput Ig and TCR repertoire sequencing analysis in rhesus macaques

**DOI:** 10.1101/2021.08.17.456682

**Authors:** Evan S. Walsh, Tammy S. Tollison, Hayden N. Brochu, Brian I. Shaw, Kayleigh R. Diveley, Hsuan Chou, Lynn Law, Allan D. Kirk, Michael Gale, Xinxia Peng

## Abstract

Recent advancements in microfluidics and high-throughput sequencing technologies have enabled recovery of paired heavy- and light-chains of immunoglobulins (Ig) and VDJ- and VJ-chains of T cell receptors (TCR) from thousands of single cells simultaneously in humans and mice. Despite rhesus macaques being one of the most well-studied model organisms for the human adaptive immune response, high-throughput single cell immune repertoire sequencing assays are not yet available due to the complexity of these polyclonal receptors. Here we employed custom primers that capture all known rhesus macaque Ig and TCR isotypes and chains that are fully compatible with a commercial solution for single cell immune repertoire profiling. Using these rhesus specific assays, we sequenced Ig and TCR repertoires in over 60,000 cells from cryopreserved rhesus PBMC, splenocytes, and FACS-sorted B and T cells. We were able to recover every Ig isotype and TCR chain, measure clonal expansion in proliferating T cells, and pair Ig and TCR repertoires with gene expression profiles of the same single cells. Our results establish the ability to perform high-throughput immune repertoire analysis in rhesus macaques at the single cell level.

## Introduction

Immunoglobulin (Ig) and T-cell receptor (TCR) repertoire analysis plays a key role in understanding the development of host immunity. These receptor molecules are responsible for recognizing a myriad of foreign antigens from infectious agents. The ability of T and B lymphocytes to give rise to such a diversity of receptor molecules with affinity to these potential antigens is, in part, due to their generation and structure. Igs are tetrameric proteins typically composed of two identical light chains (IgL or IgK) and two identical heavy chains (IgH).^1^ TCRs are heterodimeric proteins composed of paired beta (TCRβ) and alpha (TCRα) or gamma (TCRγ) and delta (TCRδ) chains, respectively. ^2^ The IgH chain, as well as TCRβ and TCRδ chains, consist of Variable (V), Diversity (D), Joining (J), and Constant (C) region gene segments. Ig light chains, as well as TCRα and TCRγ chains, do not possess D gene segments. Germline V(D)J gene segments exist at large loci within the genome and are somatically rearranged to produce functional and diversified mRNA transcripts and proteins. It is estimated that, within a single individual, the number of possible TCR and Ig V region domains is on the order of 10^13^ and 10^17^, respectfully.^3^

Increasingly, high-throughput single cell-based sequencing techniques are being employed to profile Ig and TCR repertoires. Single cell immune repertoire sequencing (scIRS) is a promising, new sequencing approach that allows for paired V(D)J repertoire analysis of thousands of cells simultaneously. For example, it has been used to identify multiple neutralizing antibodies against SARS-Cov-2 infection in humans.^4,5^ However, scIRS assays are species specific and the development of scIRS assays relies on the complete Ig and TCR reference sequences for the species of interest. Ig and TCR loci are characterized by high levels of repetitive sequences and allelic variation, making targeted sequencing and assembly difficult technical challenges.^6^ To our knowledge, current commercial scIRS assays are only available for human and mouse.

Due to the close phylogenetic relationship and highly similar physiology to humans, rhesus macaques (*Macaca mulatta*) have been one of the most popular and well-studied nonhuman primates (NHPs) for modeling immune responses in humans. ^7,8^ For example, rhesus macaques have been used to model the adaptive immune response and progression of infectious diseases from such agents varicella zoster^9^, HIV^10- 12^, and SARS-Cov-2^13^, as well as many other immune-related studies and diseases such as allograft rejection^14^ and graft versus host disease (GvHD)^15^. Recently, using long read transcriptome sequencing, we generated the first complete reference set of constant regions of all known isotypes and chain types of rhesus Ig and TCR repertoires.^16^ We also designed *in silico* rhesus-specific scIRS assays that remove the need for primers conventionally targeting variable regions.

In this study, we sought to experimentally validate and optimize rhesus specific scIRS assays that are fully compatible with commercial solutions for single cell immune repertoire profiling. Based on the complete rhesus macaque constant region reference set of Ig and TCR isotypes and chains^16^, we designed and validated primers that target these constant regions in mRNA transcripts. We further adopted these rhesus-specific primers into the human single cell immune profiling workflow provided by 10x Genomics. These rhesus-specific scIRS assays were validated using cryopreserved PBMC and splenocytes as well as FACS-sorted B and T cells from various rhesus animals. We were able to recover every known Ig and TCR isotype and pair Ig/TCR repertoire analysis with transcriptome profiles from the same single cells. We also observed clonal expansion in proliferating versus non-proliferating rhesus T cells. These results establish the ability to perform high throughput scIRS analysis in rhesus macaques with comparable performances to commercially available platforms.

## Methods

### Allogeneic mixed lymphocyte reaction (MLR) and FACS gating strategies

PBMC were obtained from peripheral blood of rhesus macaques using CPT tubes with Sodium Citrate (BD Biosciences, San Jose, CA). Cells from spleens of rhesus macaques were obtained via manual maceration of the spleen, and lysis of red blood cells using high yield lysing solution (Life Technologies, Carlsbad, CA). Cells were cryopreserved in 10% dimethyl-sulfoxide (Sigma-Aldrich, Raleigh, NC) and 90% fetal bovine serum (Corning, Corning, NY) using temperature controlled freezing containers. To prepare the mixed lymphocyte reactions (MLRs), cryopreserved cells were obtained, thawed, and counted. CD14+ monocytes were isolated from splenocytes using magnetically labeled antibodies (Miltinyi, Auburn, CA). These cells were cultured in R10 Media [Consisting of RPMI 1640 supplemented with Pen-Strep at 100U/mL, L-Glutamine at 2mM (all Gibco, Gaithersburg, MD) and 10% FCS] with IL-4 (Miltinyi Biotech, Auburn, CA) and GM-CSF (Miltinyi Biotech, Auburn, CA) for 7 total days in with the addition of TNF-α (Novus Biologicals, Centennial, CO) on the 6^th^ day to induce activated dendritic cells (DC). PBMC were thawed and labeled using Violet Proliferation Dye-450 (BD Biosciences, San Jose, CA) and combined with DCs at a 1:4 ratio. These cells were co-cultured in R10 for 5 days. At this time, the culture was stained with CD3-APC Cy7 (BD Biosciences, San Jose, CA, 557757) and sent for cell sorting on a BD DIVA (BD Biosciences, San Jose, CA). Our gating strategy is shown in Supplemental Figure 1.

### Rhesus samples, single cell processing, and cDNA generation

Multiple sample types from rhesus macaques were used to optimize and validate rhesus specific scIRS assays: cryopreserved PBMCs and splenocytes, FACS-sorted stimulated B-cells, and FACS-sorted stimulated non-proliferating and proliferating T-cells. Cells were washed in resuspension buffer (RPMI, 10% FBS), collected by centrifugation at 700xg, and resuspended at an appropriate concentration, between 700 and 1200 cells/ul. All cells were counted by hemocytometer and assayed for viability using a Countess II automated cell counter (Thermo Fisher Scientific, Waltham, MA). From each sample a maximum volume containing 17,000 cells, having approximately 90% of viable cells, were loaded onto the 10x Chromium (10x Genomics, Pleasanton, CA) controller for a targeted cell recovery of 10,000 cells (10x Genomics, Pleasanton, CA, Chromium Next GEM Single Cell V(D)J Reagents Kits v1.1_RevE). Two replicates of the same PBMC sample were established (PMBC1 and PMBC2); whereas, only one sample was prepped for each of the remaining cell types. cDNA amplification was carried out at 13 cycles of amplification for all samples and assayed for quality and concentration using a Bioanalyzer 2100 and High Sensitivity DNA Kit (Agilent, Santa Clara, CA).

### Rhesus V(D)J and 5’GEX library construction and sequencing

Construction of V(D)J libraries required the implementation of rhesus-specific primers^16^ for two subsequent enrichments of Ig and TCR transcripts as is typical for the 10x V(D)J workflow (Supplemental Table I). qPCR results of various primer pools, which varied primer concentrations to enrich for lower abundant transcripts, showed no impact on final sequencing data. Therefore, an equimolar ratio of gene-specific primers was used to construct each pool, whereby Ig assays utilized a final concentration of 0.5uM of each gene specific primer paired with 1uM 10x forward primer, and TCR assays utilized a final concentration of 1um of each gene specific primer paired with 2uM of the 10x forward primer. Due to initial sequencing results, later Ig and TCR primer pools were split into VDJ and t chain pools to improve transcript capture and chain pairing efficiency. Primer pools were constructed in volumes of 10uL, requiring the nuclease-free water volume added to the cDNA at the beginning of target enrichment to be reduced to 28ul from 33uL. Other than the needed changes to accommodate novel primers to the 10x VDJ workflow, VDJ and 5’GEX libraries were prepared according to the manufacturer’s instructions (10x Genomics, Pleasanton, CA, Chromium Next GEM Single Cell VDJ Reagents Kits v1.1).

Final V(D)J and 5’ GEX libraries were run on a Bioanalyzer 2100 with a High Sensitivity DNA kit (Agilent, Santa Clara, CA) to assess library size and concentration. Further analysis of library concentration was performed using a Qubit 3 fluorometer (Thermo Fisher Scientific, Waltham, MA) coupled with the sizing data from the Bioanalyzer to determine appropriate loading concentration for each library. Sequencing was performed using a NextSeq 500 sequencer (Illumina, San Diego, CA) utilizing 150 cycle kits in a paired end fashion. Runs were programmed to generate a 26bp read 1 sequence consisting of the 16bp 10x barcode and 10bp 10x UMI, an 8bp index read, and a 133bp read 2 sequence of the cDNA insert to exhaust the number of cycles available in the kit. Sequencing depth was targeted at a minimum of 5,000 reads per cell for V(D)J libraries and 20,000 reads per cell for 5’ GEX libraries as recommended by the 10x workflow (10x Genomics, Pleasanton, CA, Chromium Next GEM Single Cell VDJ Reagents Kits v1.1).

### Single cell V(D)J sequencing data analysis

Raw sequencing data were demultiplexed and converted to fastq files also using the Cellranger (v4.0.0) command mkfastq. Next, raw reads were de novo assembled into contigs using the Cellranger vdj command. A de novo, rather than reference-based, contig assembly was performed due incomplete annotation of the rhesus germline V(D)J sequences. To annotate the constant regions of the assembled V(D)J transcripts, assembled contigs were searched against inner-enrichment primers and a reference of constant region amplification sequences using the ublast (e-value ≤ 1e-4) and usearch_global (sequence ID threshold = 0.9) commands respectively from the USEARCH (v10.0.240) ^17^ suite of sequence analysis tools. Subsequently, to annotate the V regions of the assembled V(D)J transcripts, the assembled contigs were aligned to a custom IgBLAST (v1.8.0) ^18^ database of human and rhesus germline V(D)J sequences. An e-value cutoff of 0.01 and default parameters were used in IgBLAST queries. After annotation, we systematically filtered assembled contigs from our analysis as follows. Contigs representing misassembled chimeras (e.g. IgL primer alignment and IgM constant region alignment) or off-target transcripts (e.g. constant region primer match at the 3’ end but without a corresponding constant region alignment) were removed. Further, assembled contigs without a productive and complete V(D)J sequence or that had a premature stop codon were removed. Only contigs that passed all of these quality control criteria were considered in downstream analyses, including the assessment of pairing efficiency.

We assessed read coverage of filtered contigs using the all_contig.bam file outputted by cellranger vdj, where sequencing reads were aligned to contigs after assembly. Reads were only aligned to contigs of their respective cell barcode. For each dataset, the samtools depth command was run on the bam file to generate the read coverage per contig base position. To facilitate the comparison of read coverages across the contigs of different lengths, the read coverages were normalized as follows. For each assembled contig, we first labeled each base position as part of the 5’UTR, V region, or C region, based on the start positions of the V gene segments and the end positions of the J gene segments from IgBLAST results. Base positions within each labeled region were placed into 100 bins of equal length and the mean relative coverage per bin was calculated.

VDJ versus VJ chain pairing efficiency was assessed using B and T cells defined by the 5’ GEX data collected from the same cell samples described below. After removing low-quality cells and identifying the B and T cell clusters, the remaining cell barcodes were used as a ground truth to assess chain pairing efficiency in V(D)J sequence data since the same cell barcodes were already independently validated as a productive B or T lymphocyte based on the filtered V(D)J transcript contigs. The V(D)J sequences of each cell were integrated into a Seurat object as metadata for gene expression and clonotype analysis.

Expanded clones were defined by more than one cell sharing the same V germline gene segments and with at least 85% identical CDR3 nucleotides sequences, for both VDJ and VJ contigs. Cells that had unpaired contigs were not considered in the clonotyping analysis. CDR3 sequence clustering was performed using the CD-HIT (v4.6.6)^19^ sequence analysis tool.

### Single cell 5’ gene expression profile analysis

Raw sequencing data were demultiplexed, aligned to the RheMac10 genome annotation, and UMI-collapsed using the 10x Genomics cellranger (v4.0.0) commands mkfastq and count, respectively. Raw gene expression matrices were normalized and scaled using the SCTransform^20^ method with the Seurat R package (v3.1.5). Quality control was performed on each dataset to remove poor quality cells. For each sample, cells that had greater than 5% of their UMIs map to mitochondrial genes were removed from the analysis.

Principal component analysis was performed using the normalized and scaled expression levels of the 3,000 most variable genes in each dataset. Based on the Seurat’s recommendations, the first 30 principal components were used as input to UMAP dimension reduction and K-nearest neighbor cell clustering. Canonical marker gene expression and differential expression testing were used to determine the cell types present in the tissue culture samples. Differential expression analysis was performed using a Wilcoxon rank sum test within the FindMarkers Seurat function.

The potential for CD3+ B cells to be technical B+T cell doublets was assessed by simulating B+T cell doublets by combining UMI counts from randomly selected pairs of B and T cells. The counts for gene j in doublet i with parent cells a and b was defined as y_i,j_ = x_a,j_ + x_b,j_. The new simulated doublets were added to the unnormalized gene expression matrix while the original pair of singlets was removed. Next, the standard Seurat analysis workflow was performed as described as above. Once cells were embedded with individual UMAP coordinates the Euclidian distance between CD3+ B cells and simulated doublets was measured. This analysis was performed using between 10 and 100 simulated doublets by an increase of 10 cells. The simulation and distance calculations were repeated 10 times for each number of simulated doublets. Wilcoxon rank sum tests were performed to test for a difference in distance between CD3+ B cells to each other and to simulated B+T cell doublets.

## Results

### High throughput single cell Ig and TCR sequencing in rhesus macaque

As shown in Fig. 1, we adopted rhesus specific V(D)J primers into the human single cell immune profiling assays commercially available from 10x Genomics and sequenced in total over 80,000 single cell barcodes from cryopreserved rhesus PBMC, splenocytes, FACS-sorted memory B cells (CD20+ CD27+) and stimulated T cells (CD3+; sorted into proliferating and non-proliferating fractions by dye dilution). De novo assembly of raw sequencing reads resulted in over 150,000 unfiltered, unannotated rhesus V(D)J transcript contigs. We devised a custom computational pipeline for contig isotype and germline annotation, as well as quality control filtering standards (see Methods). Assembled VDJ and VJ transcript sequences were searched against a custom IgBLAST database of rhesus and human germline segments and selected for sequences encoding complete, productive variable domains. Filtering using these criteria yielded the final set of 63,623 (78.8%) cell barcodes with a total of 97,932 (61.8%) V(D)J sequences from six rhesus samples (two PBMCs, splenocytes, sorted memory B cells, proliferating and non-proliferating T cells) (Supplemental Table II).

**Fig 1.**
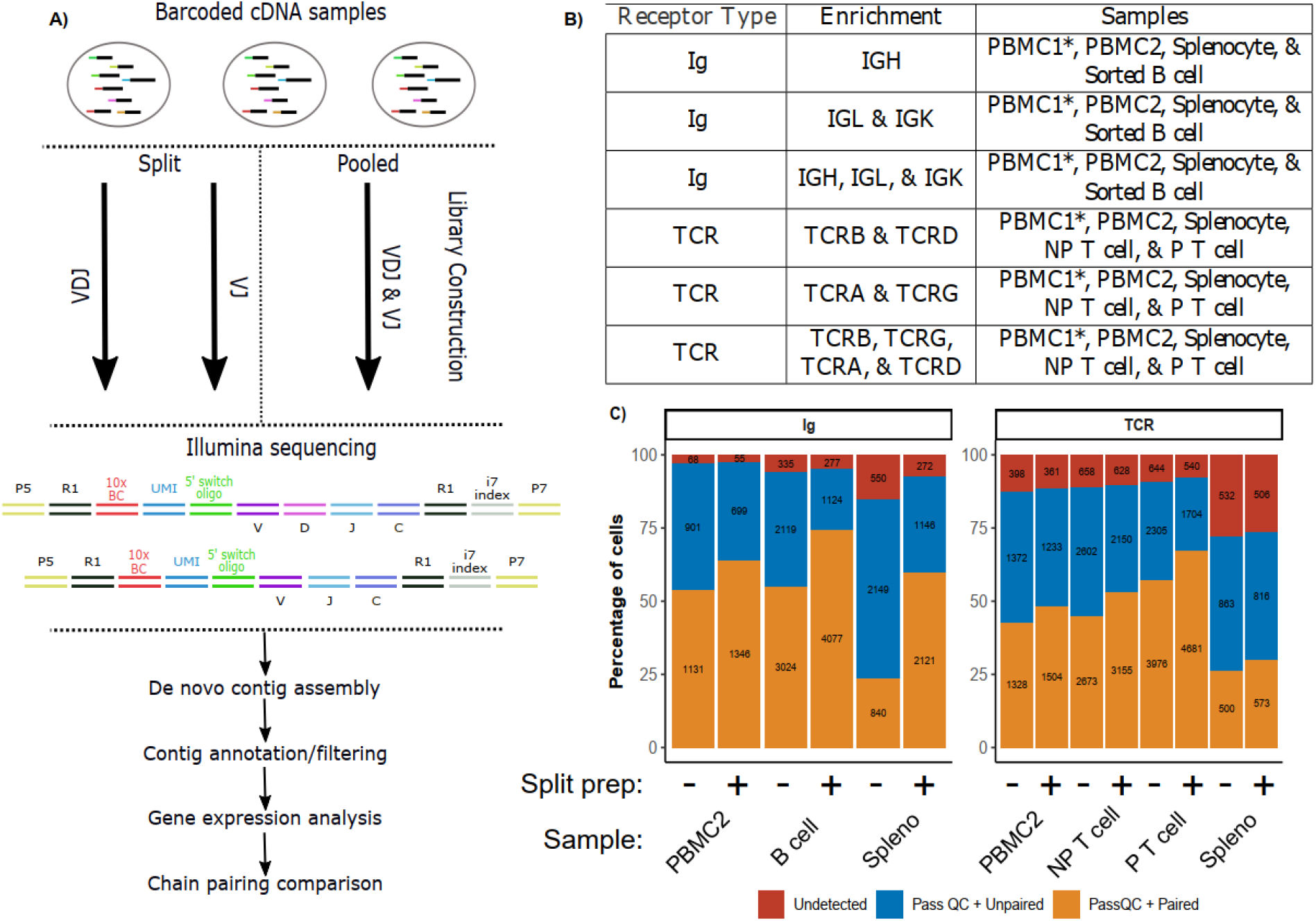
Overview of assay development. a) Droplet-based scRNA-seq was used to generate an initial barcoded cDNA libraries. To maximize the rate at which cells with paired VDJ and VJ contigs were recovered, sequencing libraries were generated using pooled and separate primers for VDJ and VJ chain types. After sequencing and data analysis the chain pairing rate was assessed and compared between the two configurations using filtered B and T cell barcodes recovered from 5’ gene expression analysis. b) Targeted enrichment performed on each cDNA sample. The PBMC1 sample is labeled with a * to indicate that a gene expression library was not constructed for this dataset, so a comparison between split and pooled library preps was not performed. c) Bar charts indicate the percentage of cell barcodes with either no V(D)J contigs that pass quality control (QC) filtering, an unpaired contig(s) that does pass, or paired contigs that do pass. In each sample, the fraction of paired contigs that pass QC increases when VDJ and VJ primers are split in separate library preps.

Despite the 5’ capture method used in the 10x immune profiling assays; we did not observe significant bias in read coverage toward the 5’ end of assembled V(D)J transcript contigs. Instead, we observed that the read coverages were normally distributed across the 5’ untranslated regions (UTRs) and variable regions of the obtained V(D)J contigs and were uniformly distributed across the constant region (Supplemental Fig. 2a). We reasoned that coverage was distributed differently in the constant region due to the coincidence that the read 2 length (133 bp) was similar to the lengths of the 5’ portion of constant regions captured by our assays. The median length of constant region sequences amplified from our enrichment protocol was between 73-108 bp (Supplemental Figure 2b). Therefore, those sequencing reads corresponding to the unfragmented cDNA molecules would cover the entire amplified constant region. The average numbers of reads supporting the final filtered contigs in Ig enrichment libraries were between 117 and 2,434 and the numbers of unique molecular identifiers (UMI) were between 1 and 9. The median numbers of reads and UMIs supporting the final filtered contigs in TCR enrichment libraries were comparable, ranging from 763 to 2,282 for reads and from 1 to 4 for UMIs (Supplemental Table II).

To improve the recovery of paired VDJ and VJ sequences from the same cells (e.g. TCRβ and TCRα or IgG and IgL chains) we devised a sequencing library construction strategy in which VDJ and VJ transcripts were enriched in both separate and pooled sequencing libraries (Fig. 1a-b). Pairing efficiency, defined as the percentage of cells with at least one final filtered VDJ and one final filtered VJ sequence, may be affected by differences in VDJ- and VJ-chain transcript abundances as well as individual primer amplification efficiencies. To facilitate the comparisons of pairing efficiency between VDJ and VJ split and pooled libraries, we additionally sequenced and generated gene expression profiles of the same single cells from five rhesus samples. B and T cell barcodes that passed gene expression quality control filtering served as an independent “ground-truth” of viable B and T cells for assessing pairing efficiency. When VDJ and VJ primers were used for separate library preparation, an increase in the fraction of cells with paired VDJ-VJ repertoires was observed across all five samples and both Ig and TCR enrichments (Fig. 1c). Between 60 and 74% of filtered B cell barcodes from gene expression libraries had paired heavy and light chains (Supplemental Table III). We observed between 30 and 68% of T cell barcodes that had paired VDJ and VJ sequences (Supplemental Table III). Both Ig and TCR pairing efficiency ranges were comparable to other scIRS assays in the split library configurations.^21,22^ We noted a relatively lower pairing efficiency rate observed in T cells from the one splenocyte sample. Overall, we recovered a total of 7,539 B cells and 9,906 T cells with paired VDJ and VJ contigs as well as a filtered gene expression profile (Supplemental Tables II & III).

### B cell receptor repertoire profiling in rhesus PBMC, splenocyte, and sorted B cell samples

We analyzed BCR repertoires from two separate PBMC samples (the total number of B cells sequenced n=3,418 and n=3,035), one splenocyte (n=15,682), and one sorted B cell sample (n= 9,746) (Supplemental Table II). The number of cell barcodes with at least one filtered V(D)J sequence will be higher in the absence of paired GEX data due to the inability to filter out doublets, non-viable cells, or ambient transcripts assigned to cell barcodes. Overall, we detected every known Ig isotype (Fig. 2a) and were also able to recover every known IgA allotype and IgG subclass identified in our previous Iso-Seq analysis, observing preferential usage of IgA*01 and IgA*02 as well as IgG1, respectively.^16^ No IgE contigs were detected in the PBMC2 and splenocyte samples, however IgE is known to be detected at the lowest levels among any IgH isotype.^22^ Ig light chains (IgK and IgL) were more frequently detected than IgH contigs across all four samples. We also investigated the ratio of cells with IgA, IgE, and IgG contigs to those with IgM and IgD. IgM and IgD isotypes are known to be expressed in mature naive B cells while IgA, IgE, and IgG are expressed by activated (memory) B cells after undergoing an IgH class-switch recombination process. ^23^ Overall, this IgH ratio ranged between 0.39 to 0.60 for our four Ig datasets, suggesting that we might have captured about twice as many naive relative to activated B cells, on average (Supplemental Fig. 2d). As expected, the largest ratio of activated to naïve B cells appeared in the FACS B cell dataset, which were sorted using cell surface markers of B cell memory (CD20 & CD27).

**Fig 2.**
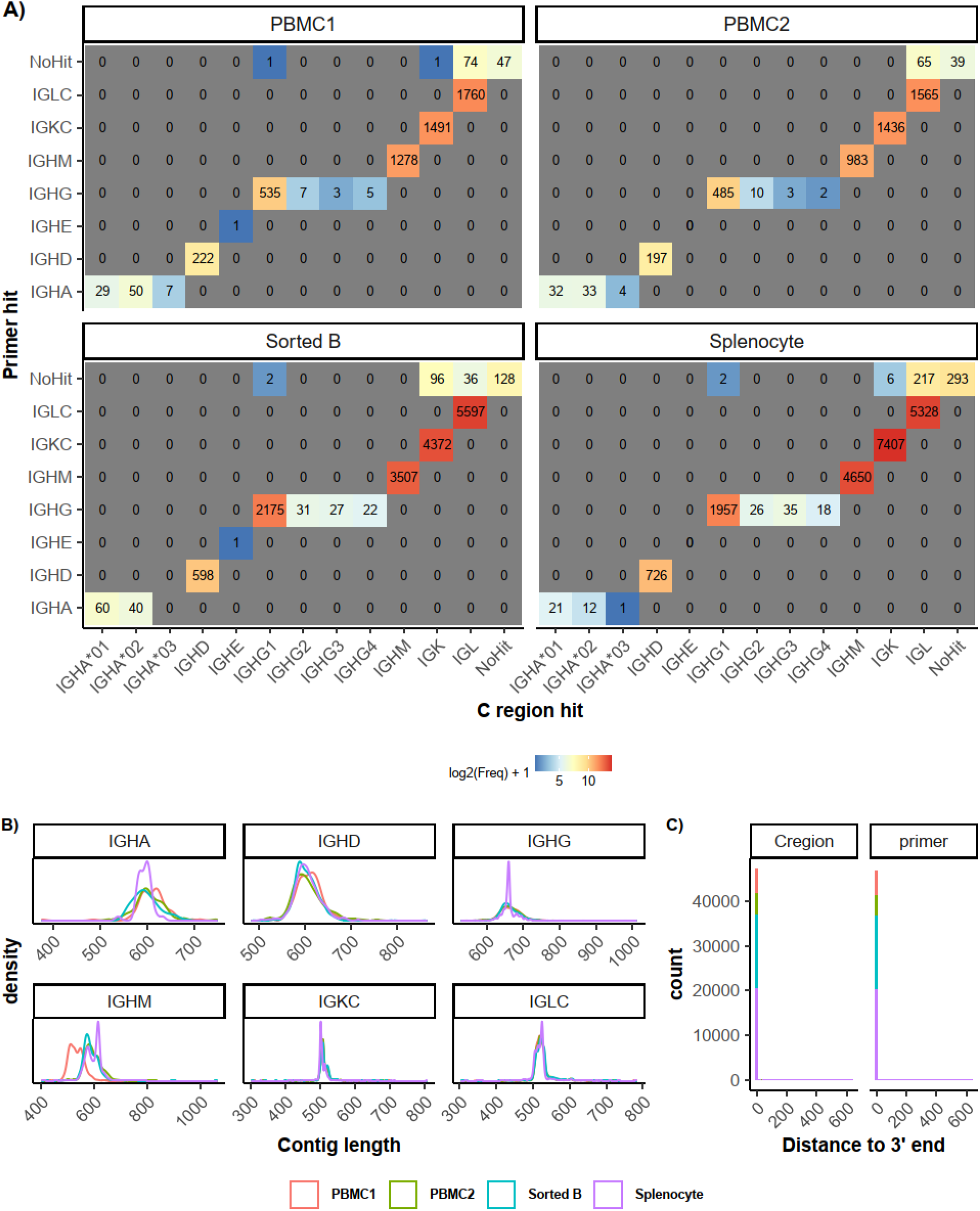
Characterization of isotypes Ig enrichment libraries. a) Recovery of 3’ of enriched V(D)J transcripts. Tiled plots indicate the number of contigs with respective primer and constant region alignments. b) Distribution of contig length by Ig isotype across the Ig enrichment datasets. IGHE was omitted because of the low number of contigs detected. A different IgM inner enrichment primer was used for the PBMC1 library construction which was 59 base pairs 5’ of the primer used for all other datasets. This is consistent with the differences in the length distributions. c) Distribution of the distances between constant region and primer alignments and 3’ end of contigs.

Despite the 5’ capture technology used to build the cDNA libraries, we observed a high rate of complete assembly of the constant regions at the 3’ end of the obtained Ig V(D)J contigs. Only 1.6 to 2.5% of filtered Ig V(D)J contigs did not map to an inner enrichment primer or expected amplified constant region sequence (Fig. 2a). Ig V(D)J contigs that failed to map to the inner enrichment primer or amplified constant region sequences also had a lower sequencing depth in terms of both reads and UMIs (Supplementary Fig. 1c). To ensure that the 3’ constant regions of Ig V(D)J sequences were correctly assembled we examined the distances of our primer and constant region alignments from the 3’ end of Ig V(D)J contigs. We found that 97.4-98.4% map directly to the 3’ end of Ig V(D)J contigs (Fig. 2c). The distribution of isotype contig lengths was also largely consistent across the different datasets, providing confidence that our assembled contigs represent true B cell receptor mRNA molecules (Fig 2b).

### T cell receptor repertoire profiling in rhesus PBMC, splenocyte, and sorted T cell samples

Similarly, we analyzed T cell receptor repertoires from two separate PBMC samples (the total number of T cells sequenced n=3,988, n=3,543), one splenocyte (n=3,162), and two samples derived from a MLR experiment representing proliferating (P, n= 12,964) and non-proliferating (NP, n= 8,085) FACS-sorted T cells (Supplemental Table II). Like our B cell repertoire analysis, TCR V(D)J contigs were searched against the inner-enrichment primers used to build TCR libraries in addition to a reference of expected amplified constant region sequences. Importantly, in addition to the more commonly targeted α- and β-TCRs, we also included primers capturing δ- and γ-TCRs. Notably, primers that target δ- and γ-chains are not currently available in the human and mouse single cell immune profiling assays from 10x Genomics, resulting in repertoire analyses solely covering αβ T cells. ^25,26^ γδ T cells have been reported to make up only 4% of T cells, on average. ^27^ As expected, we detected δ- and γ-chain types at a lower frequency than the predominantly expressed α- and β-chain types (Fig. 3a). However, we observed TCRα+ cells to be only as much as 1.59 times more abundant than TCRγ+ cells and TCRβ+ cells were about 15.8 times more abundant than TCRδ+ cells in mixed cell datasets (i.e PBMC and splenocytes), suggesting that leaving out γ- and δ-TCRs could potentially miss relevant biological insights. Interestingly, overall TCR VDJ contigs were 1.38x more abundant than VJ contigs in αβ T cells, but 4.52x less abundant in γδ T cells (Fig 3a).

**Fig 3.**
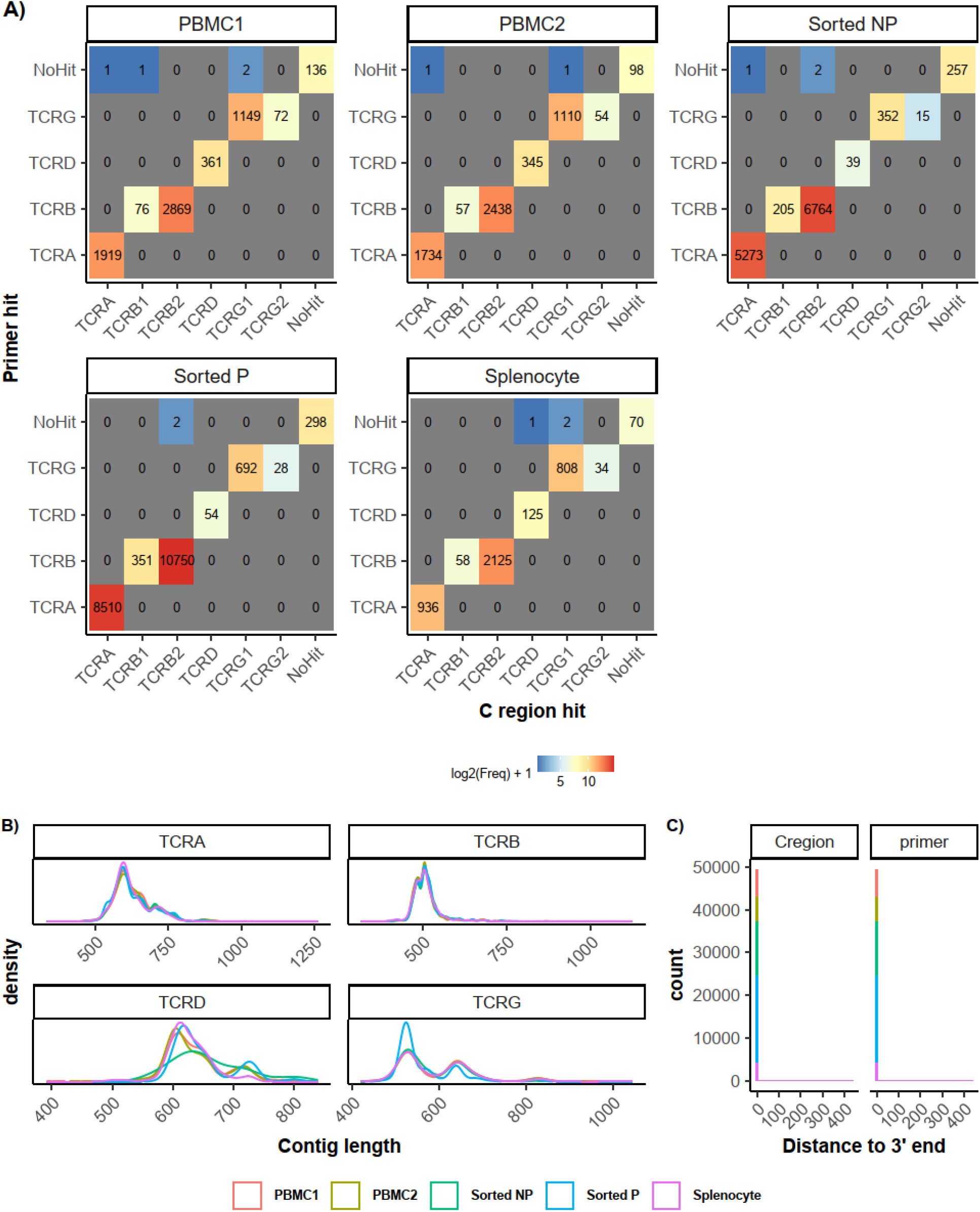
Characterization of isotypes in TCR enrichment libraries. a) Recovery of 3’ of enriched V(D)J transcripts. Tiled plots indicate the number of contigs with respective primer and constant region alignments. b) Distribution of contig length by TCR isotype across the TCR enrichment datasets. c) Distribution of the distances between constant region and primer alignments and 3’ end of contigs.

Similar to our Ig repertoire analysis, we also observed a high rate of complete constant region assembly in our TCR enrichment datasets. Only 1.5 to 2.2% of TCR V(D)J contigs did not map to an inner enrichment primer or amplified constant region sequence (Fig. 3a). Furthermore, TCR V(D)J contigs that failed to map to the inner enrichment primer sequences or constant region sequences also had a lower sequencing depth in terms of reads and UMIs than those that did map to our constant region and primer references (Supplemental Fig. 2c). Correct constant region assembly was confirmed by observing constant region amplification sequences and inner-enrichment primers mapped directly to the 3’ end of the TCR V(D)J contigs, between 97.8-98.5% (Fig. 3c). The distributions of TCR V(D)J contig lengths were also largely consistent across different samples, supporting that our assembled TCR contigs represented the full-length TCR V(D)J transcripts (Fig. 3b).

### Clonal expansion in proliferating T cells

To assess the potential of our rhesus scIRS assays to detect clonally expanded lineages, we compared the clonal expansion rates in our proliferating (P) and non-proliferating (NP) T cell datasets. These cells were isolated from a MLR, in which responding PBMCs were labeled with a proliferation dye and incubated with MHC mismatched activated DCs for five days using a previously described protocol (Supplemental Fig. 1).^28^ We then sorted these cells based on CD3 positivity and whether they had undergone proliferation as indicated by a loss of proliferation dye. Proliferating cells were enriched for those cells responding to alloantigens from the MHC mismatched DCs. We defined T-cell lineages by grouping cells with identical VDJ and VJ germline variable gene segments and required at least 85% nucleotide identity in both CDR3 regions, although the number of clones did not change significantly when this sequence identity threshold was increased (Fig. 4a).

**Fig 4.**
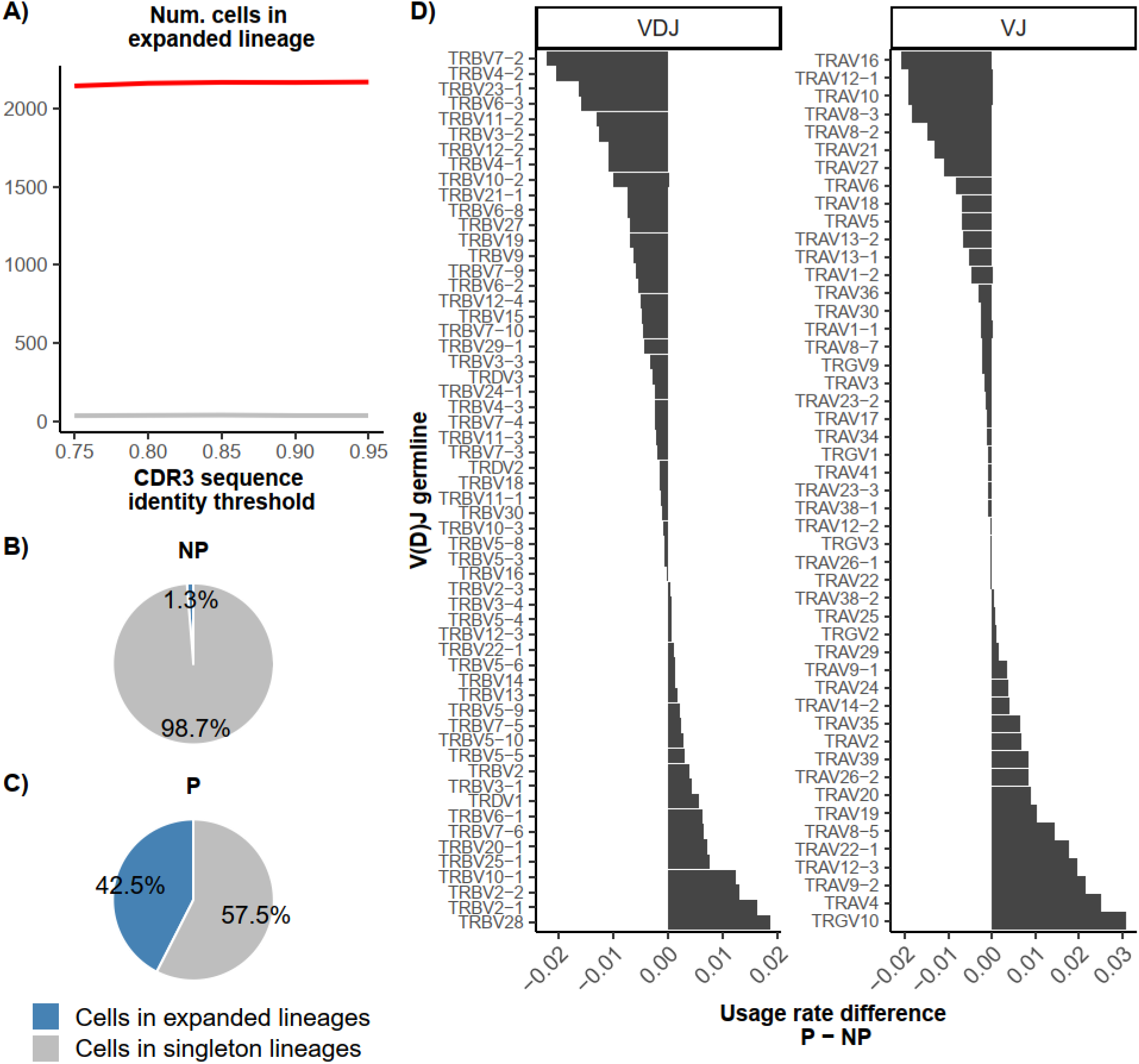
Detection of clonally expanded lineages in proliferating T cells. a) Number of cells in expanded lineages in NP and P T cells as a function of the CDR3 sequence identity threshold. b) Lineage expansions among proliferating (P) sorted T cells. Pie charts indicate percentage of cells belonging to expanded lineages. c) Same as b), but for non-proliferating (NP) T cells. d) Usage rate differences of variable germline gene segments between clonally expanded P T cells and singleton NP T cells. Usage rate is defined as the number of cells a germline gene is present in relative to the total number of cells.

As expected, we observed a substantial clonal expansion in the proliferating T cell dataset with 42.5% of cells belonging to clonally expanded lineages compared to < 2% in the non-proliferating T cell dataset (Fig 4a-c). Despite the high clonal expansion rate in P T cells, we did not observe any lineages that dominated the clonally expanded cells. Out of the 2,166 cells of expanded lineages in the P T cell dataset, 530 (24.5%) belonged to a lineage of size two. The largest lineage size consisted of only 24 cells. Hierarchical clustering of TCR VDJ and VJ variable germline gene segments in clonally expanded P T cells did not reveal any modules of highly overlapping VJ and VDJ combinations (Supplemental Fig. 3a). MLRs typically result in a generalized activation of some T cells predisposed for proliferation. There are a multitude of potential antigens that can activate T cells, there is a propensity of stimulatory cytokines to induce by-stander proliferation in previously primed cells (e.g. those with a memory phenotype), and heterologous cross-reactive stimulation can be expected.^29^ Nevertheless, the products of an MLR are substantially enriched for allospecific cells.

Interestingly, among P T cells the most frequent VJ germline gene segment detected in expanded lineages was TRGV10, which was co-detected in many T cells with TCRβ molecules (Supplemental Fig. 3a&c and Supplemental Fig. 4). Additionally, TCRGV10 showed the largest increase in usage between expanded P T cells and NP T cells among any VJ or VDJ germline segment (Fig 4d). This observation was intriguing, given the high correlation of germline usage rate between expanded P and NP T cells (Supplemental Fig. 3b). In some of the clonotypes, no additional TCRβ contig was detected, but the frequency of αβ T cells that also expressed a γ chain showed the largest increase from NP to expanded P T cells (Supplemental Fig. 4). Similar to what others have reported, we observed a diverse number of TCRα and TCRβ chains and a limited number of TCRγ chains in our γ-αβ T cells in both NP and P + expanded T cell datasets.^30^ In both NP and P + expanded γ-αβ T cells the TCRGV10 variable gene was used in over 80% of lineages (singleton or expanded). We did not observe one TCRα and TCRβ variable gene used in over 12% of lineages.

Some studies have reported T cells that co-express αβ and γδ receptors. ^30,31^ Others have characterized δ/αβ T cells which possess chimeric contigs that encompass a δ variable gene but α joining and constant region domains.^32^ This chimeric scenario is unlikely to be the case in this data because chimeric contigs as the one described in δ/αβ T cells would be removed from our analyses (see Methods). The frequency of γ-αβ T cells was between 9.32%-12.46% of the T cells in PBMC and splenotye samples, while γ-β T cells was between 11.77%-25.39%. It shows that the observed co-expression was not unique to these FACS-sorted T cell datasets. Since typical single cell based TCR repertoire sequencing analyses have focused on αβ T cells, it is unclear if these αβ T cells co-expressing TCRγ or TCRδ tend to be undetected or are atypically deleted at some point post-production. Additionally, the presence of TCRγ transcripts in αβ T cells does not imply γδ TCRs are on the surface of the same T cells. Obviously future analyses of more samples are needed. However, this complete chain pairing coverage highlights the unbiased potential of our rhesus scIRS assays which could also enable future exploratory analysis of αβ and γδ co-expressions.

### Integrative analysis of gene expression profiles and V(D)J repertoires of the same single cells

Parallel 5’ gene expression profiling analysis allowed us to cluster the same single cells into groups of cell types, and to overlay individual cells with their Ig and TCR repertoire data (Fig. 5a-b). For example, in the PBMC2 sample (Fig. 5a), we identified 2,100 B cells across 3 cell clusters. We recovered 1,344 (64.1%) B cells with paired heavy and light chains, 636 (30.3%) with only a light chain, and 62 (3%) with only a heavy chain. Among the 3,071 T cells in the same PBMC2 sample, we recovered 1,495 (48.7%) cells with paired TCR VDJ and VJ chains, 651 (21.2%) cells with only a VDJ and 582 (19%) cells with only a VJ chain. The rates of paired TCR chains were similar across the different T cell subtypes. Among the 3,531 B cells identified in the splenocyte sample (Fig. 5b), we recovered 2,118 (60%) B cells with paired heavy and light chains, 817 (23%) with only a light chain, 328 (9.3%) with only a heavy chain. In the same splenocyte sample we captured 1,886 T cells. Of those, we recovered 575 (30.5%) cells with paired TCR VDJ and VJ chains, 472 (25%) cells with only a VDJ, and 347 (18.4%) cells with only a VJ chain. In addition, we observed 492 (26.1%) T cells without any TCR V(D)J contigs (Supplemental Table III).

**Fig 5.**
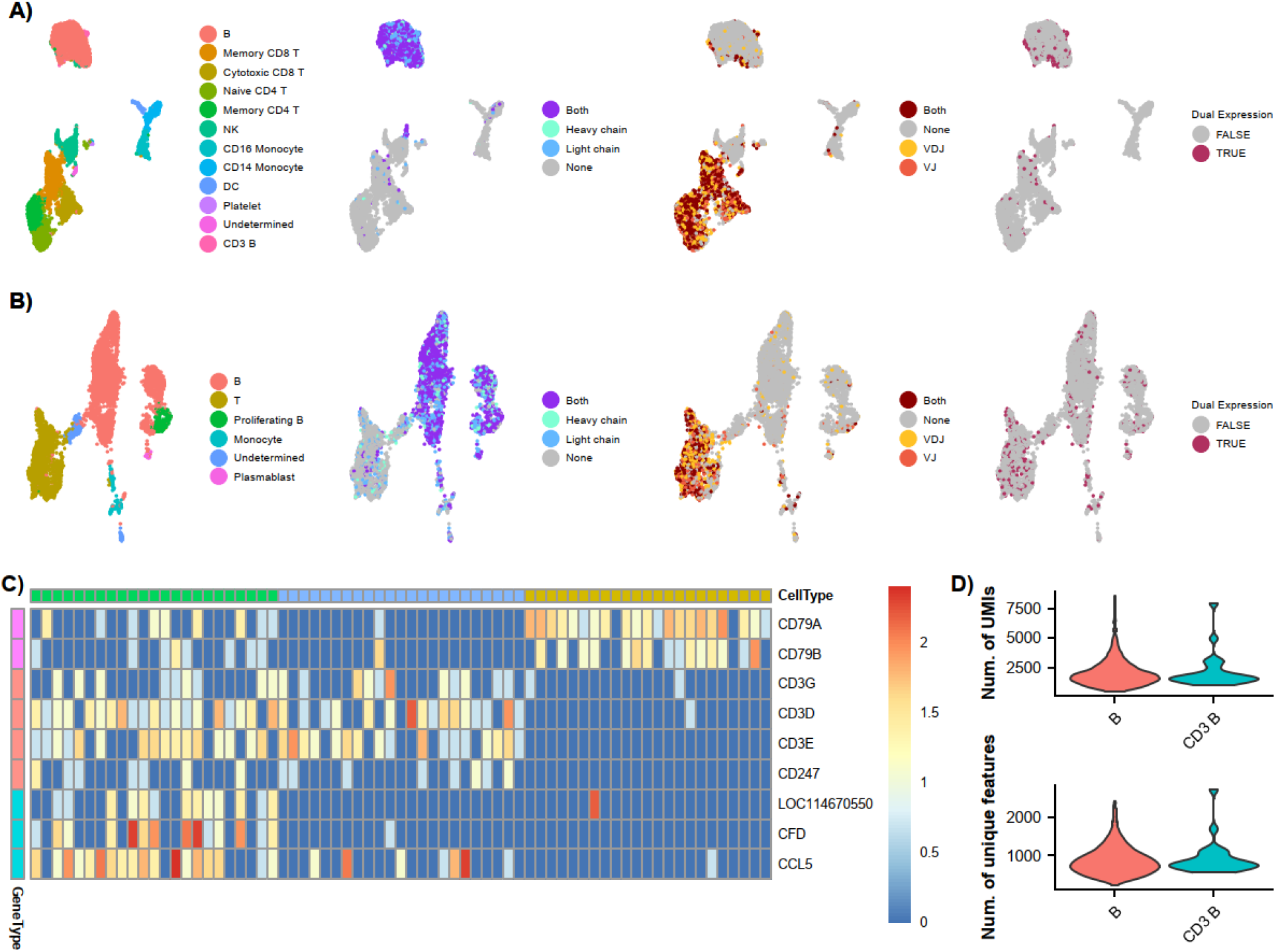
Integration of gene expression and V(D)J profiles of single cells. a) UMAP plots of the (left to right) assignment of cell type, BCR chains, TCR chains, and dual BCR and TCR detection for each cell in PBMC 2 sample. b) Same as a) but the splenocyte sample. c) Normalized expression of Ig and TCR signal transduction genes, as well as CD3+ B cell gene expression signatures across CD3 B cells (green) and randomly sampled B (gold) and T cells (blue). d) Distribution of UMIs and unique features in CD3+ B and B cells.

Intriguingly, among the cells sequenced in the PBMC2 dataset we observed a small cluster of 23 B cells clustered separately from the others (Fig 5a). Upon investigation, we obtained both Ig and TCR V(D)J contigs from a high fraction of these cells. Of these 23 cells, Ig and TCR dual expression was detected in 22 (> 95%) cells. Dual expression of Ig and TCR molecules was also observed in 229 additional cells, but these cells appeared to be randomly distributed in UMAP space. Therefore, we focused our exploratory Ig/TCR dual expression analysis on this small cluster of 23 cells.

Dual expression of Ig and TCRs or other B and T cell markers have been reported by others, but the role that these hybrid lymphocytes play is largely unknown. ^33-35^ The possibility of doublets, where two cells are given the same cellular barcode, does arise when using droplet-based single cell technologies such as the 10x Genomics used here. To assess the potential for these cells to be B+T cell doublets we used differential expression analysis to identify transcriptomic signatures that were absent in B and T cells. This small B cell clusters not only expressed both Ig and TCR molecules, but the genes involved in B and T cell receptor signal transduction. CD79A, CD79B, CD3E, CD3G, and CD3D were all upregulated in th is small cluster of B cells. As such we labeled this cluster as CD3+ B cells. The top differentially expressed genes appeared to be constitutively expressed across these CD3+ B cells while absent in regular B and T cells (Fig. 5c). Furthermore, CD3+ B cells did not have a greater sequencing depth than other B cells in terms of number of UMIs or unique features expressed (Fig 5d). To further demonstrate the transcriptional differences of CD3+ B cells from technical B+T cell doublets we simulated B+T cell doublets and added their expression profiles to the gene expression matrix to rerun dimension reduction and clustering (see Methods). To assess the transcriptional similarity of CD3+ B cells and simulated B+T cell doublets we measured the Euclidean distance between CD3+ B cells to each other and to simulated doublets. Consistently, CD3+ B cells were closer to each other (3.06 ± 0.397) (i.e more transcriptionally similar) than to simulated doublets (5.14 ± 0.570). These results were additionally found to be statistically significant by Wilcoxon rank sum test (adjusted p-value < 0.05). Taken together, these data suggest that these CD3+ B cells were less likely to be B and T cell doublets, but more likely to be expressing both Ig and TCR transcripts.

## Discussion

In this study, using extensive experimental optimization and validation, we established rhesus specific single cell high-throughput assays for Ig and TCR repertoire sequencing analysis that recover every known rhesus Ig and TCR isotype and chain type. While rhesus macaque is one of the most widely used NHP models, it is our understnading that this is the first such assay developed for rhesus macaques that are also fully compatible with a commercially available platform. We also devised a custom computational pipeline to process and analyze the generated rhesus single cell V(D)J sequencing data.

The establishment of these single cell-based Ig/TCR repertoire sequencing assays will improve our understanding of immune responses in rhesus macaques, which will in turn allow for applications to further understanding the immune response in humans. For example, we were able to detect clonally expanded lineages of T cells in a proliferating T cell sample, using the full-length productive VDJ and VJ sequences paired within single cells. The overlay of cellular gene expression profiles allowed the identification of the underlying cell types and subsets. We also observed rare cells expressing both Ig and TCR V(D)J transcripts that could be investigated in the future. Compared to the human and mouse assays commercially available from 10x Genomics, our assays also capture TCRγ and TCRδ chains. While αβ T cells are more frequent in general, our results showed that the frequencies of γδ T cells could be extremely variable, and sometimes their frequency could be as high as 26% of V(D)J contigs detected in a PBMC sample. This suggests that the additional coverage of γ- and δ-TCRs may enable the discovery of relevant biological insights that are overlooked by current assays.

Unexpectedly, we detected γ-αβ T cells in all four types of rhesus samples we analyzed in this study: PBMC, splenocyte, FACS sorted non-proliferating and clonally-expanded, proliferating T cell samples. Interestingly, we also observed a large increase in the frequencies of γ-αβ T cells in the clonally-expanded, proliferating T cell sample compared to the non-proliferating T cell sample (Supplemental Figure 4). Generally, a mutually exclusive pattern of γδ- and αβ-expression is expected.^36^ However, reports have demostrated that there are exceptions. Using FACS in combination with qPCR and sc-RNAseq reseraches have observed and validated the presence of dual expressing γδ-αβ T cells at the mRNA and cell-surface receptor level in multiple tissues and developmental stages of human and mice. ^30,31^ Furthermore, transgenic mice constructed with a pair of productively rearranged γ and δ genes have been shown to produce a normal number of αβ T cells^37^, in addition to other reported unconventional expression patterns of TCR molecules.^38,39^ Considering potential techincal confounding factors such as ambinet transcripts and techincal doublets, future independent analyses will be needed to experimentally investigate the rhesus γ-αβ T cells observed here.

Analysis of the adaptive immune response has co-evolved with the advancement of high-throughput IRS strategies. For RNA-based analysis, 5’ RACE has been used to reliably generate unbiased amplification of Ig and TCR cDNA molecules and can be done using a simple primer set. ^40-42^ The major limitations of 5’ RACE and similar strategies has been extracting the pairing of variable region information required for the determination of antigen/epitope specificity. Therefore, to recover paired variable region sequences, single cell high-throughput sequencing strategies started to emerge. Emulsion PCR, where TCRβ and TCRα RNA molecules from single cells are fused together^43,44^ and pairSeq where single cells are isolated and mRNA reverse transcription reactions occur individually within a 96-well plate became commonplace. ^45^ Despite the advancement in chain pairing of these IRS strategies, these methods are generally low-throughput in terms of the number of cells sequenced in a single experiment and are not applicable to many Ig and TCR chain types. The recent advent of droplet-based single cell sequencing technologies has greatly improved the throughput of IRS analysis. The human and mouse immune profiling assays from 10x Genomics allow for the recovery of paired, full-length V(D)J sequences as well as gene expression profiles of thousands of cells simultaneously.

However, the development of similar scIRS assays for other species is challenging. For a species of interest, it requires the complete Ig and TCR reference sequences, which is often lacking due to the extreme complexity of Ig and TCR loci. Recently, we obtained a complete reference set for the constant regions of rhesus Ig and TCR isotypes and chain types using long read transcriptome sequencing^16^, which bypasses the need of rhesus specific germline reference sequences. This reference set enabled us to design and validate primers targeting rhesus V(D)J constant region sequences to achieve complete and unbiased repertoire analysis, whereas primer pools targeting the highly diverse variable regions can miss as much as 50% of repertoire sequences.^15^ Here, we used de novo assembly to reconstruct captured Ig/TCR transcripts with full-length productive V(D)J regions instead of using germline reference sequence based approaches. Still, the availability of comprehensive rhesus germline reference sequences could greatly facilitate the analysis and the biological interpretation of the obtained Ig/TCR repertoire data. Building rhesus germline reference sequence collections would require future developments.

In summary, these results demonstrate that scIRS assays we established here for rhesus macaques can be used to recover every known Ig and TCR isotype and chain type in an unbiased fashion, pair VDJ and VJ transcripts in the same cells, detect clonally expanded lineages and dual expressing cels, and integrate V(D)J repertoires with cellular gene expression profiles. These scIRS assays will greatly facilitate the analysis of the adaptive immune responses in rhesus macaques.

## Acknowledgments

Cryopreserved splenocyte samples were provided by Dr. Sallie R. Permar at Duke University. Rhesus B cell line BLCL was provided by the NIH Nonhuman Primate Reagent Resource. Rhesus T cell lines RH447 and RH444 were provided by Dr. Vanessa M. Hirsch at NIAID/NIH.

## Footnotes

This project has been funded in part with Federal funds from the National Institute of Allergy and Infectious Diseases, National Institutes of Health, Department of Health and Human Services, under Contract No. HHSN272201800008C, and by the Washington National Primate Research Center Core grant P51 OD010425 from the National Institutes of Health, Office of the Director (MG), and grant R21AI120713 (to X.P.).

The nonhuman primate work has also been funded by multiple awards from the National Institute for Allergy and Infectious Diseases including R38AI40297 (supporting B.I.S.) and U19AI131471 (to A.D.K.).

## Abbreviations used in this article

NHP: nonhuman primate
GvHD: graft versus host disease
IRS: Immune repertoire sequencing
RACE: Rapid Amplification of cDNA ends
MPCR: multiplex PCR
IMGT: the international ImMunoGeneTics information system
Iso-Seq: full-length isoform sequencing
scIRS: single cell immune repertoire sequencing
UMI: unique molecular identifier
NP: non-proliferating
P: proliferating
UTR: untranslated region
DC: dendritic cell

5’ gene expression sequencing data presented in this article have been submitted to the National Center for Biotechnology Information GEO under the accession number GSE179722. Datasets generated during the current study are available at the NCBI Sequence Read Archive under project number PRJNA746267.

## References

1. Schroeder, H. W., & Cavacini, L. (2010). Structure and Function of Immunoglobulins. The Journal of Allergy and Clinical Immunology, 125(2 0 2), S41–S52. 10.1016/j.jaci.2009.09.046

2. Davis, M. M., & Bjorkman, P. J. (1988). T-cell antigen receptor genes and T-cell recognition. Nature, 334(6181), 395–402. 10.1038/334395a0

3. Charles A Janeway, J., Travers, P., Walport, M., & Shlomchik, M. J. (2001). The Generation of Lymphocyte Antigen Receptors. Immunobiology: The Immune System in Health and Disease. 5th Edition, 2001. https://www.ncbi.nlm.nih.gov/books/NBK10774/. Accessed Apr 12, 2021.

4. Li, F., Luo, M., Zhou, W., Li, J., Jin, X., Xu, Z., Juan, L., Zhang, Z., Li, Y., Liu, R., Li, Y., Xu, C., Ma, K., Cao, H., Wang, J., Wang, P., Bu, Z., & Jiang, Q. (2020). Single cell RNA and immune repertoire profiling of COVID-19 patients reveal novel neutralizing antibody. Protein & Cell, 10.1007/s13238-020-00807-6

5. Mor, M., Werbner, M., Alter, J., Safra, M., Chomsky, E., Lee, J. C., Hada-Neeman, S., Polonsky, K., Nowell, C. J., Clark, A. E., Roitburd-Berman, A., Ben-Shalom, N., Navon, M., Rafael, D., Sharim, H., Kiner, E., Griffis, E. R., Gershoni, J. M., Kobiler, O., … Freund, N. T. (2021). Multi-clonal SARS-CoV-2 neutralization by antibodies isolated from severe COVID-19 convalescent donors. PLoS Pathogens, 17(2), e1009165. 10.1371/journal.ppat.1009165

6. Watson, C. T., Glanville, J., & Marasco, W. A. (2017). The Individual and Population Genetics of Antibody Immunity. Trends in Immunology, 38(7), 459–470. 10.1016/j.it.2017.04.003

7. Williams, K., Lackner, A., & Mallard, J. (2016). Non-human primate models of SIV infection and CNS neuropathology. Current Opinion in Virology, 19, 92–98. 10.1016/j.coviro.2016.07.012

8. Flynn, J. L., Gideon, H. P., Mattila, J. T., & L in, P. L. (2015). Immunology studies in non-human primate models of tuberculosis. Immunological Reviews, 264(1), 60–73. 10.1111/imr.12258

9. Meyer, C., Kerns, A., Haberthur, K., Dewane, J., Walker, J., Gray, W., & Messaoudi, I. (2013). Attenuation of the adaptive immune response in rhesus macaques infected with simian varicella virus lacking open reading frame 61. Journal of Virology, 87(4), 2151–2163. 10.1128/JVI.02369-12

10. Lackner, A. A., & Veazey, R. S. (2007). Current concepts in AIDS pathogenesis: insights from the SIV/macaque model. Annual Review of Medicine, 58, 461–476. 10.1146/annurev.med.58.082405.094316.

11. Hansen, S. G., Ford, J. C., Lewis, M. S., Ventura, A. B., Hughes, C. M., Coyne-Johnson, L., Whizin, N., Oswald, K., Shoemaker, R., Swanson, T., Legasse, A. W., Chiuchiolo, M. J., Parks, C. L., Axthelm, M. K., Nelson, J. A., Jarvis, M. A., Piatak, M., Lifson, J. D., & Picker, L. J. (2011). Profound early control of highly pathogenic SIV by an effector memory T-cell vaccine. Nature, 473(7348), 523–527. 10.1038/nature10003

12. Hansen, S. G., Piatak, M., Ventura, A. B., Hughes, C. M., Gilbride, R. M., Ford, J. C., Oswald, K., Shoemaker, R., Li, Y., Lewis, M. S., Gilliam, A. N., Xu, G., Whizin, N., Burwitz, B. J., Planer, S. L., Turner, J. M., Legasse, A. W., Axthelm, M. K., Nelson, J. A., … Picker, L. J. (2013). Immune clearance of highly pathogenic SIV infection. Nature, 502(7469), 100–104. 10.1038/nature12519

13. Munster, V. J., Feldmann, F., Williamson, B. N., van Doremalen, N., Pérez-Pérez, L., Schulz, J., Meade-White, K., Okumura, A., Callison, J., Brumbaugh, B., Avanzato, V. A., Rosenke, R., Hanley, P. W., Saturday, G., Scott, D., Fischer, E. R., & de Wit, E. (2020). Respiratory disease in rhesus macaques inoculated with SARS-CoV-2. Nature, 585(7824), 268–272. 10.1038/s41586-020-2324-7

14. Knechtle, S. J., Shaw, J. M., Hering, B. J., Kraemer, K., & Madsen, J. C. (2019). Translational impact of NIH-funded nonhuman primate research in transplantation. Science Translational Medicine, 11(500) 10.1126/scitranslmed.aau0143

15. Miller, W. P., Srinivasan, S., Panoskaltsis-Mortari, A., Singh, K., Sen, S., Hamby, K., Deane, T., Stempora, L., Beus, J., Turner, A., Wheeler, C., Anderson, D. C., Sharma, P., Garcia, A., Strobert, E., Elder, E., Crocker, I., Crenshaw, T., Penedo, M. C. T., … Kean, L. S. (2010). GVHD after haploidentical transplantation: a novel, MHC-defined rhesus macaque model identifies CD28-CD8+ T cells as a reservoir of breakthrough T-cell proliferation during costimulation blockade and sirolimus-based immunosuppression. Blood, 116(24), 5403–5418. 10.1182/blood-2010-06-289272

16. Brochu, H. N., Tseng, E., Smith, E., Thomas, M. J., Jones, A. M., Diveley, K. R., Law, L., Hansen, S. G., Picker, L. J., Gale, M., & Peng, X. (2020). Systematic Profiling of Full-Length Ig and TCR Repertoire Diversity in Rhesus Macaque through Long Read Transcriptome Sequencing. Journal of Immunology (Baltimore, Md.: 1950), 204(12), 3434–3444. 10.4049/jimmunol.1901256

17. Edgar, R. C. (2010). Search and clustering orders of magnitude faster than BLAST. Bioinformatics (Oxford, England), 26(19), 2460–2461. 10.1093/bioinformatics/btq461

18. Ye, J., N. Ma, T. L. Madden, and J. M. Ostell. 2013. IgBLAST: an immunoglobulin variable domain sequence analysis tool. Nucleic Acids Res. 41(W1): W34–W40. 10.1093/nar/gkt382

19. Fu, L., Niu, B., Zhu, Z., Wu, S., & Li, W. (2012). CD-HIT: accelerated for clustering the next-generation sequencing data. Bioinformatics (Oxford, England), 28(23), 3150–3152. 10.1093/bioinformatics/bts565

20. Hafemeister, C., & Satija, R. (2019). Normalization and variance stabilization of single cell RNA-seq data using regularized negative binomial regression. Genome Biology, 20(1), 296. 10.1186/s13059-019-1874-1

21. Goldstein, L. D., Chen, Y. J., Wu, J., Chaudhuri, S., Hsiao, Y., Schneider, K., Hoi, K. H., Lin, Z., Guerrero, S., Jaiswal, B. S., Stinson, J., Antony, A., Pahuja, K. B., Seshasayee, D., Modrusan, Z., Hötzel, I., & Seshagiri, S. (2019). Massively parallel single cell B-cell receptor sequencing enables rapid discovery of diverse antigen-reactive antibodies. Communications Biology, 2(1), 1–10. 10.1038/s42003-019-0551-y

22. Singh, M., Al-Eryani, G., Carswell, S., Ferguson, J. M., Blackburn, J., Barton, K., Roden, D., Luciani, F., Giang Phan, T., Junankar, S., Jackson, K., Goodnow, C. C., Smith, M. A., & Swarbrick, A. (2019). High-throughput targeted long-read single cell sequencing reveals the clonal and transcriptional landscape of lymphocytes. Nature Communications, 10(1), 3120. 10.1038/s41467-019-11049-4

23. Jr, Janeway, C. A., Travers, P., Walport, M., & Shlomchik, M. J. (2001). Immunobiology (5th ed.). Garland Science.

24. Tong, P., Granato, A., Zuo, T., Chaudhary, N., Zuiani, A., Han, S. S., Donthula, R., Shrestha, A., Sen, D., Magee, J. M., Gallagher, M. P., van der Poel, Cees E., Carroll, M. C., & Wesemann, D. R. (2017). IgH isotype-specific B cel l receptor expression influences B cell fate. Proceedings of the National Academy of Sciences of the United States of America, 114(40), E8411–E8420. 10.1073/pnas.1704962114

25. De Simone, M., Rossetti, G., & Pagani, M. (2018). Single Cell T Cell Receptor Sequencing: Techniques and Future Challenges. Frontiers in Immunology, 910.3389/fimmu.2018.01638

26. Carter, J. A., Preall, J. B., Grigaityte, K., Goldfless, S. J., Jeffery, E., Briggs, A. W., Vigneault, F., & Atwal, G. S. (2019). Single T Cell Sequencing Demonstrates the Functional Role of αβ TCR Pairing in Cell Lineage and Antigen Specificity. Frontiers in Immunology, 1010.3389/fimmu.2019.01516

27. Chien, Y., Meyer, C., & Bonneville, M. (2014). γδ T cells: first line of defense and beyond. Annual Review of Immunology, 32, 121–155. 10.1146/annurev-immunol-032713-120216

28. Espinosa, J., Herr, F., Tharp, G., Bosinger, S., Song, M., Farris, A. B., George, R., Cheeseman, J., Stempora, L., Townsend, R., Durrbach, A., & Kirk, A. D. (2016). CD57(+) CD4 T Cells Underlie Belatacept-Resistant Allograft Rejection. American Journal of Transplantation: Official Journal of the American Society of Transplantation and the American Society of Transplant Surgeons, 16(4), 1102–1112. 10.1111/ajt.13613

29. DeWolf, S., Grinshpun, B., Savage, T., Lau, S. P., Obradovic, A., Shonts, B., Yang, S., Morris, H., Zuber, J., Winchester, R., Sykes, M., & Shen, Y. (2018). Quantifying size and diversity of the human T cell alloresponse. JCI Insight, 3(15) 10.1172/jci.insight.121256

30. Edwards, S. C., Sutton, C. E., Ladell, K., Grant, E. J., McLaren, J. E., Roche, F., Dash, P., Apiwattanakul, N., Awad, W., Miners, K. L., Lalor, S. J., Ribot, J. C., Baik, S., Moran, B., McGinley, A., Pivorunas, V., Dowding, L., Macoritto, M., Paez-Cortez, J., … Mills, K. H. G. (2020). A population of proinflammatory T cells coexpresses αβ and γδ T cell receptors in mice and humans. The Journal of Experimental Medicine, 217(5) 10.1084/jem.20190834

31. Reitermaier, R., Krausgruber, T., Fortelny, N., Ayub, T., Vieyra-Garcia, P. A., Kienzl, P., Wolf, P., Scharrer, A., Fiala, C., Kölz, M., Hiess, M., Vierhapper, M., Schuster, C., Spittler, A., Worda, C., Weninger, W., Bock, C., Eppel, W., & Elbe-Bürger, A. (2021). αβγδ T cells play a vital role in fetal human skin development and immunity. Journal of Experimental Medicine, 218(e20201189) 10.1084/jem.20201189

32. Pellicci, D. G., Uldrich, A. P., Le Nours, J., Ross, F., Chabrol, E., Eckle, S. B. G., de Boer, R., Lim, R. T., McPherson, K., Besra, G., Howell, A. R., Moretta, L., McCluskey, J., Heemskerk, M. H. M., Gras, S., Rossjohn, J., & Godfrey, D. I. (2014). The molecular bases of δ/αβ T cell–mediated antigen recognition. The Journal of Experimental Medicine, 211(13), 2599–2615. 10.1084/jem.20141764

33. Japp, A. S., Meng, W., Rosenfeld, A. M., Perry, D. J., Thirawatananond, P., Bacher, R. L., Liu, C., Gardner, J. S., Atkinson, M. A., Kaestner, K. H., Brusko, T. M., Naji, A., Luning Prak, E. T., & Betts, M. R. (2021). TCR+/BCR+ dual-expressing cells and their associated public BCR clonotype are not enriched in type 1 diabetes. Cell, 184(3), 827-839.e14. 10.1016/j.cell.2020.11.035

34. Ahmed, R., Omidian, Z., Giwa, A., Cornwell, B., Majety, N., Bell, D. R., Lee, S., Zhang, H., Michels, A., Desiderio, S., Sadegh-Nasseri, S., Rabb, H., Gritsch, S., Suva, M. L., Cahan, P., Zhou, R., Jie, C., Donner, T., & Hamad, A. R. A. (2019). A Public BCR Present in a Unique Dual-Receptor-Expressing Lymphocyte from Type 1 Diabetes Patients Encodes a Potent T Cell Autoantigen. Cell, 177(6), 1583-1599.e16. 10.1016/j.cell.2019.05.007

35. Liu, Y., Ye, S., Guo, X., Li, W., Xia, Y., Wen, X., Yu, J., Jia, Y., Liu, X., Guo, Y., & Zhao, Y. (2020). Discovery and characteristics of B cell-like T cells: A potential novel tumor immune marker? Immunology Letters, 220, 44–50. 10.1016/j.imlet.2020.01.007

36. Hayes, S., Li, L., & Love, P. (2005). TCR Signal Strength Influences αβ/γδ Lineage Fate. Immunity, 10.1016/j.immuni.2005.03.014

37. Ishida, I., Verbeek, S., Bonneville, M., Itohara, S., Berns, A., & Tonegawa, S. (1990). T-cell receptor gamma delta and gamma transgenic mice suggest a role of a gamma gene silencer in the generation of alpha beta T cells. PNAS, 10.1073/pnas.87.8.3067

38. Bowen, S., Sun, P., Livak, F., Sharrow, S., & Hodes, R., (2014). A Novel T Cell Subset with Trans-Rearranged Vγ-Cβ TCRs Shows Vβ Expression Is Dispensable for Lineage Choice and MHC Restriction. Journal of Immunology, 10.4049/jimmunol.1302398

39. Hochstenbach, F., & Brenner M. (1989) T-cell receptor δ-chain can substitute for α to form a βδ heterodimer. https://doi.org/10.1038/340562a0

40. Ichinohe, T., Miyama, T., Kawase, T., Honjo, Y., Kitaura, K., Sato, H., Shin-I, T., & Suzuki, R. (2018). Next-Generation Immune Repertoire Sequencing as a Clue to Elucidate the Landscape of Immune Modulation by Host–Gut Microbiome Interactions. Frontiers in Immunology, 910.3389/fimmu.2018.00668

41. Gao, F., & Wang, K. (2015). Ligation-anchored PCR unveils immune repertoire of TCR-beta from whole blood. BMC Biotechnology, 15, 39. 10.1186/s12896-015-0153-9

42. Heather, J. M., Best, K., Oakes, T., Gray, E. R., Roe, J. K., Thomas, N., Friedman, N., Noursadeghi, M., & Chain, B. (2015). Dynamic Perturbations of the T-Cell Receptor Repertoire in Chronic HIV Infection and following Antiretroviral Therapy. Frontiers in Immunology, 6, 644. 10.3389/fimmu.2015.00644

43. Turchaninova, M. A., Britanova, O. V., Bolotin, D. A., Shugay, M., Putintseva, E. V., Staroverov, D. B., Sharonov, G., Shcherbo, D., Zvyagin, I. V., Mamedov, I. Z., Linnemann, C., Schumacher, T. N., & Chudakov, D. M. (2013). Pairing of T-cell receptor chains via emulsion PCR. European Journal of Immunology, 43(9), 2507–2515. 10.1002/eji.201343453

44. Liu, X., & Wu, J. (2018). History, applications, and challenges of immune repertoire research. Cell Biology and Toxicology, 34(6), 441–457. 10.1007/s10565-018-9426-0

45. Howie, B., Sherwood, A. M., Berkebile, A. D., Berka, J., Emerson, R. O., Williamson, D. W., Kirsch, I., Vignali, M., Rieder, M. J., Carlson, C. S., & Robins, H. S. (2015). High-throughput pairing of T cell receptor α and β sequences. Science Translational Medicine, 7(301), 301ra131. 10.1126/scitranslmed.aac5624

