## Supplemental Figures and Tables for "Single cell based high-throughput Ig and TCR repertoire sequencing analysis in rhesus macaques"

| | | Number of cells | | Number of contigs | | After filtering | | | |
|---|---|---|---|---|---|---|---|---|---|
| **Sample** | **Enrichment** | **Before filtering** | **After filtering** | **Before filtering** | **After filtering** | **UMIs** | **Reads** | **UMIs / contig** | **Reads / contig** |
| PBMC1 | Ig | 5831 | 3418 | 11374 | 5511 | 80088 | 11276276 | 14.53 | 2046.14 |
| PBMC1 | TCR | 5413 | 3988 | 11063 | 6586 | 26350 | 12386249 | 4 | 1880.69 |
| PBMC2 | Ig | 4408 | 3035 | 9690 | 4854 | 62224 | 20105960 | 12.82 | 4142.14 |
| PBMC2 | TCR | 4775 | 3543 | 9839 | 5838 | 21642 | 13991598 | 3.71 | 2396.64 |
| Sorted B cell | Ig | 11611 | 9746 | 28669 | 16692 | 233072 | 47957489 | 13.96 | 2873.08 |
| Sorted NP T cell | TCR | 9760 | 8085 | 19875 | 12908 | 57590 | 45290898 | 4.46 | 3508.75 |
| Sorted P T cell | TCR | 15279 | 12964 | 31796 | 20685 | 187013 | 48438019 | 9.04 | 2341.7 |
| Splenocyte | Ig | 18624 | 15682 | 28693 | 20699 | 330969 | 26548813 | 15.99 | 1282.61 |
| Splenocyte | TCR | 5017 | 3162 | 7580 | 4159 | 9816 | 9478924 | 2.36 | 2279.14 |

*Supplemental Table II. Summary of number of cells, contigs and sequencing depth before and after quality control filtering. Contigs representing chimeras or off-target transcripts as well as contigs without productive and in-frame V(D)J sequences were filtered from the analysis. Cells without any filtered contigs were additionally removed from the analysis. Reads and UMI support data are from the cellranger vdj all_contig_annotatinos.csv file.*

| | Enrichment | nCells_total | nCells_paired | nCells_VDJ | nCells_VJ | nCells_either | Undetected | nCells_VJ_only | nCells_VDJ_only |
|---|---|---|---|---|---|---|---|---|---|
| **PBMC2** | TCR | 3071 | 1495 (48.7%) | 2146 (69.9%) | 2077 (67.6%) | 1233 (40.1%) | 343 (11.2%) | 582 (19%) | 651 (21.2%) |
| **Splenocyte** | TCR | 1886 | 575 (30.5%) | 1047 (55.5%) | 922 (48.9%) | 819 (43.4%) | 492 (26.1%) | 347 (18.4%) | 472 (25%) |
| **Sorted_NP T cell** | TCR | 5933 | 3155 (53.2%) | 4563 (76.9%) | 3897 (65.7%) | 2150 (36.2%) | 628 (10.6%) | 742 (12.5%) | 1408 (23.7%) |
| **Sorted_P T cell** | TCR | 6925 | 4681 (67.6%) | 5799 (83.7%) | 5267 (76.1%) | 1704 (24.6%) | 540 (7.8%) | 586 (8.5%) | 1118 (16.1%) |
| **PBMC2** | Ig | 2096 | 1344 (64.1%) | 1406 (67.1%) | 1980 (94.5%) | 698 (33.3%) | 54 (2.6%) | 62 (3%) | 636 (30.3%) |
| **Splenocyte** | Ig | 3531 | 2118 (60%) | 2446 (69.3%) | 2935 (83.1%) | 1145 (32.4%) | 268 (7.6%) | 328 (9.3%) | 817 (23.1%) |
| **B cells** | Ig | 5478 | 4077 (74.4%) | 4332 (79.1%) | 4946 (90.3%) | 1124 (20.5%) | 277 (5.1%) | 255 (4.7%) | 869 (15.9%) |

*Supplemental Table III. Summary of TCR+Ig sequence recovery. The total number of cells is based on filtered gene expression analysis data. Values indicate the number of cells the with the respective V(D)J sequence. Paired: VDJ+VJ. VDJ: VDJ only + paired. VJ: VJ only + paired. Either: VJ only + VDJ only. Undetected: No VDJ or VJ.*

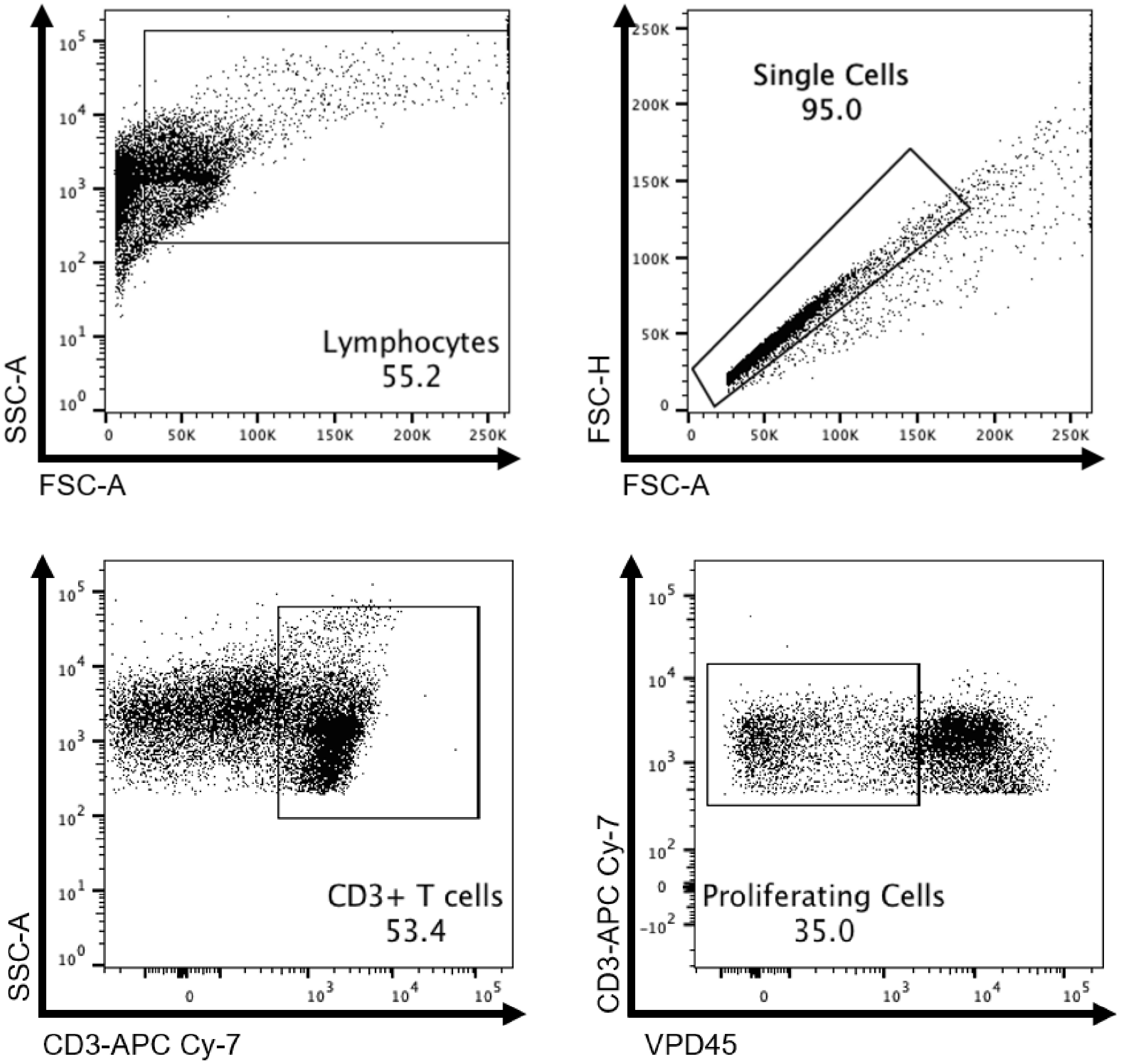


*Supplemental Figure 1. Allospecific gating strategy. Allospecific T cells were isolated by sorting size, CD3 positivity, and VPD450 proliferation dye loss.*

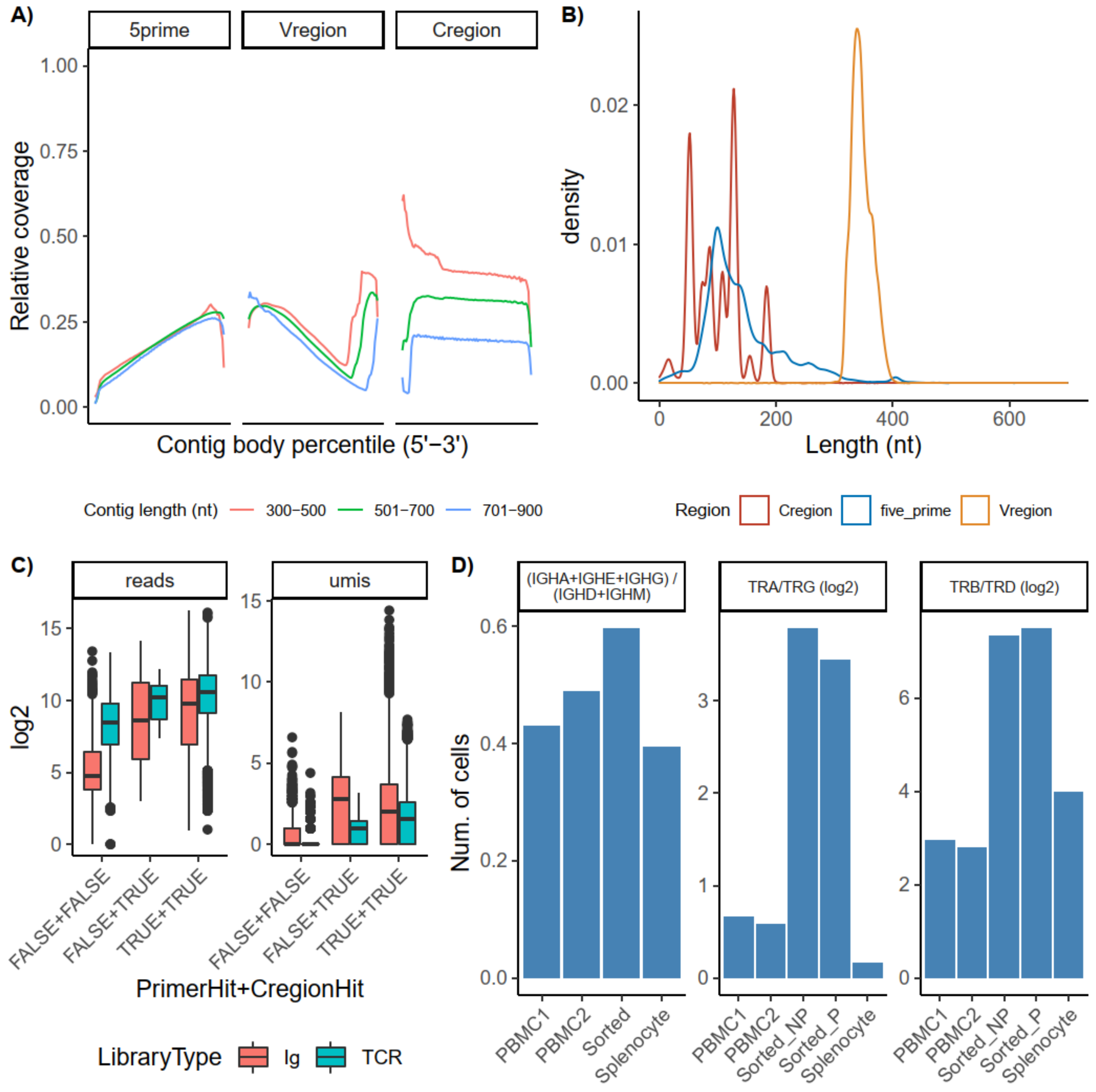


*Supplemental Figure 1. Analysis of Ig and TCR repertories. A) Distribution of contig coverage across 5' UTR, variable and constant regions of Ig and TCR enrichment datasets. The relative coverage was calculated for a contig percentile position based on how many reads mapped in that region relative to the total mapped to the contig. B) Distribution of contig length by domain. Region boundaries were determined using IgBLAST search results. C) Association between contig sequencing depth and isotype annotation. X axis labels represent whether a filtered contig maps to an amplified constant region sequence and/or an inner enrichment primer. D) Ratio of isotypes in terms of number of cells present. Left facet indicates the activated to naïve B cell ratio. Middle facet is the ratio of VDJ chain types in T cells. Right facet is the ratio of VJ chain types in T cells.*

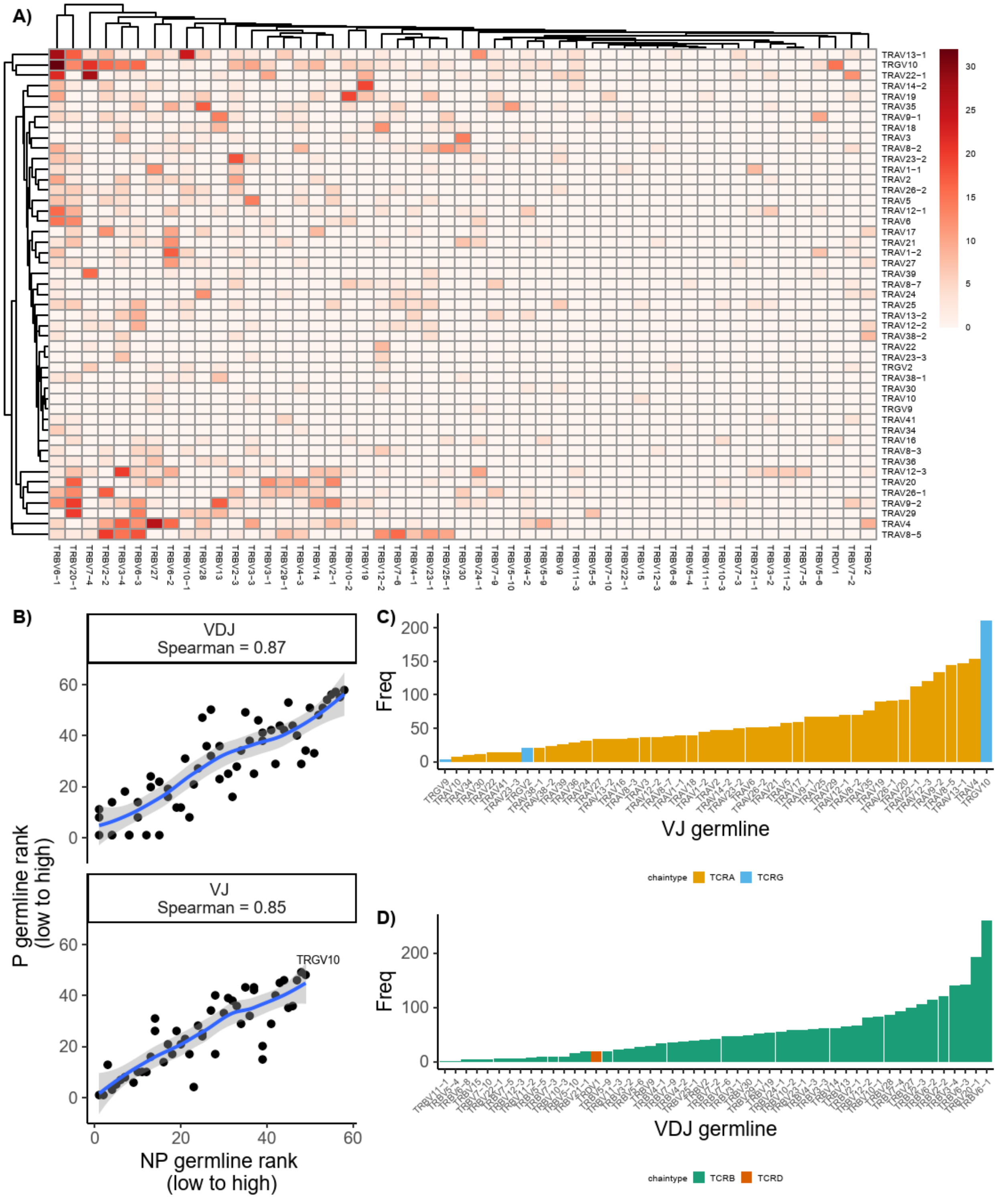


*Supplemental Figure 3. Variable region pairing and usage in clonally expanded proliferating T cells. A) Hierarchical clustering of variable VDJ (x axis) and VJ (y axis) germline gene segments. Colors indicate the number of cells with the respective chain pair. B) Spearman correlation of variable germline segments usage ranking. X axis indicates usage ranking in NP T cells. Y axis indicates ranking to expanded P T cells. C-D) Marginal densities of VDJ and VJ germline segments in expanded P T cells. Colors indicated ichain type of each variable gene.*

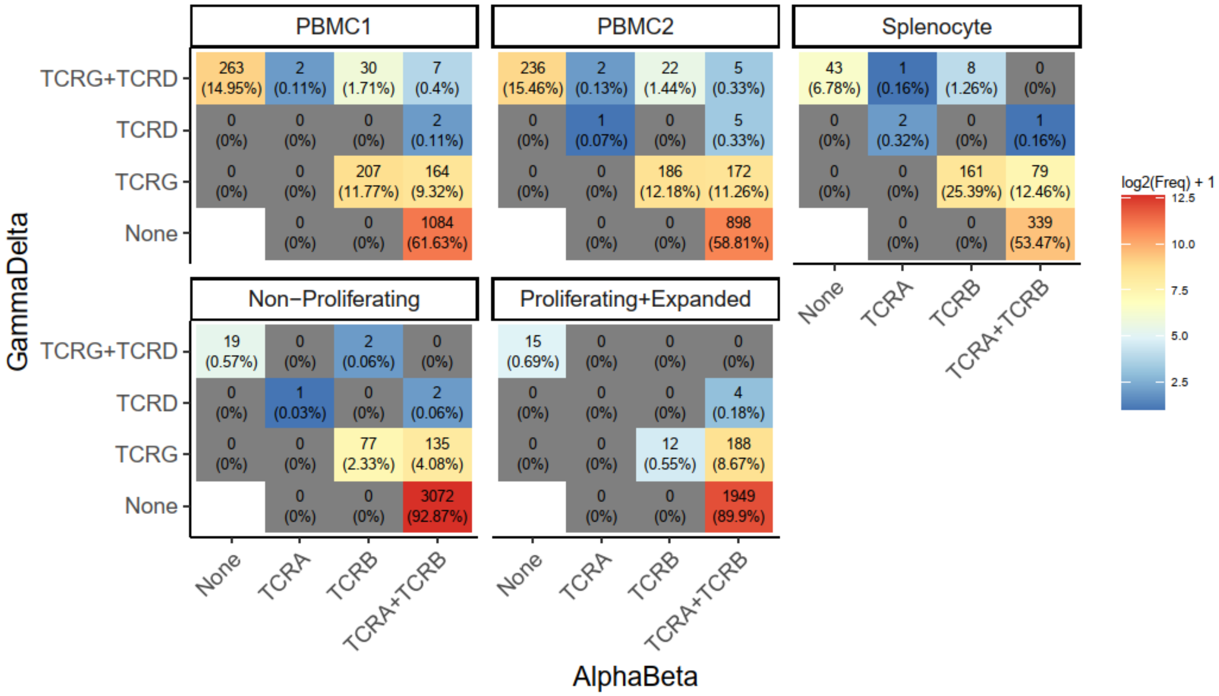


*Supplemental Figure 4. Co-expression of TCR chain types. Tiled plots indicate the number and percentage of cells with the respective TCR chain types across the five TCR enrichment datas*
